# The CUL5 ubiquitin ligase complex mediates resistance to CDK9 and MCL1 inhibitors in lung cancer cells

**DOI:** 10.1101/518555

**Authors:** Shaheen Kabir, Justin Cidado, Courtney Andersen, Cortni Dick, Pei-Chun Lin, Therese Mitros, Hong Ma, Ron Baik, Matthew Belmonte, Lisa Drew, Jacob Corn

## Abstract

Overexpression of anti-apoptotic proteins MCL1 and Bcl-xL is a frequent event in blood and solid cancers. Inhibitors targeting MCL1 are in clinical development, however many cancer models are intrinsically resistant to this approach. To discover mechanisms underlying resistance to MCL1 inhibition, we performed multiple flow-cytometry based genome-wide CRISPR screens that interrogate two drugs directly or indirectly targeting MCL1. Remarkably, both screens identified three components (CUL5, RNF7 and UBE2F) of a cullin-RING ubiquitin ligase complex (CRL5) that resensitized cells to MCL1 inhibition. We find that levels of the BH3-only pro-apoptotic proteins Bim and Noxa are proteasomally regulated by the CRL5 complex. Accumulation of Noxa caused by depletion of CRL5 components particularly skewed the balance in favor of apoptosis when cells were challenged with an MCL1 inhibitor. Discovery of a novel role of CRL5 in apoptosis and resistance to MCL1 inhibitors exposes new drug targets and the potential to improve combination treatments.

## INTRODUCTION

Cancer cells frequently manipulate the intrinsic apoptotic pathway to evade cell death and expand their proliferative capacity. Aberrant increases in levels of anti-apoptotic proteins in the BCL-2 family, such as amplification of *MCL1*, have been widely implicated in the transformation of cancer cells and the development of resistance to current therapies (Kelly & Strasser, 2011). Members of the BCL-2 family are classified based on the conservation of their BCL-2 homology (BH) domains: multi-domain proteins BAK, BAX and BOK serve as apoptosis executors in the mitochondria; proteins containing only the BH3 domain (BH3-only) promote BAK/BAX activation. Anti-apoptotic proteins such as MCL1, Bcl-xL and BCL-2 inhibit apoptosis by antagonistic binding to BH3-only pro-apoptotic proteins and BAK and BAX (Czabotar, Lessene, Strasser, & Adams, 2014).

High-resolution investigation of somatic copy number alterations has revealed gene amplification of *MCL1* and *BCL2L1* (Bcl-xL) are key determinants of survival of breast and non-small cell lung cancer (NSCLC), as depletion of MCL1 and Bcl-xL greatly induces apoptosis in cancer cells (Goodwin, Rossanese, Olejniczak, & Fesik, 2015; Xiao et al., 2015; Zhang et al., 2011). Various models of multiple myeloma, acute myeloid leukemia (AML) and B-cell acute lymphoblastic leukemia (B-ALL) are also dependent on MCL1 expression for survival (Koss et al., 2013). Amplification of *MCL1* is also a prognostic indicator for disease severity and progression (Campbell et al., 2018; Yin et al., 2016), making it an attractive therapeutic target.

In an effort to restrict the action of anti-apoptotic proteins, numerous compounds have been developed that mimic BH3-only proteins (BH3-mimetics). Unfortunately, the first BH3-mimetics that specifically antagonized Bcl-xL were associated with significant thrombocytopenia, thus complicating their therapeutic use (Lessene et al., 2013; Joel D Leverson et al., 2015; Tao et al., 2014). Small-molecule inhibition of MCL1 has recently gained significant attention (**Figure 1A**), and compounds that selectively target MCL1 are currently in clinical trials (Abulwerdi et al., 2014; Burke et al., 2015; Caenepeel et al., 2018; Kotschy et al., 2016; J D Leverson et al., 2015; S64315, n.d.; Tron et al., 2018). Promising reports of direct BH3-mimetic MCL1 inhibitors in preclinical hematological malignancies show potent efficacy with low cytotoxicity (Kotschy et al., 2016; J D Leverson et al., 2015). However, assessment of MCL1 inhibitors in solid breast tumors showed little single agent activity unless combined with a chemotherapeutic agent (Merino et al., 2017). Co-dosing Bcl-xL and MCL1 inhibitors to achieve effective treatment may be complicated by severe accompanying side effects.

**Figure 1.**
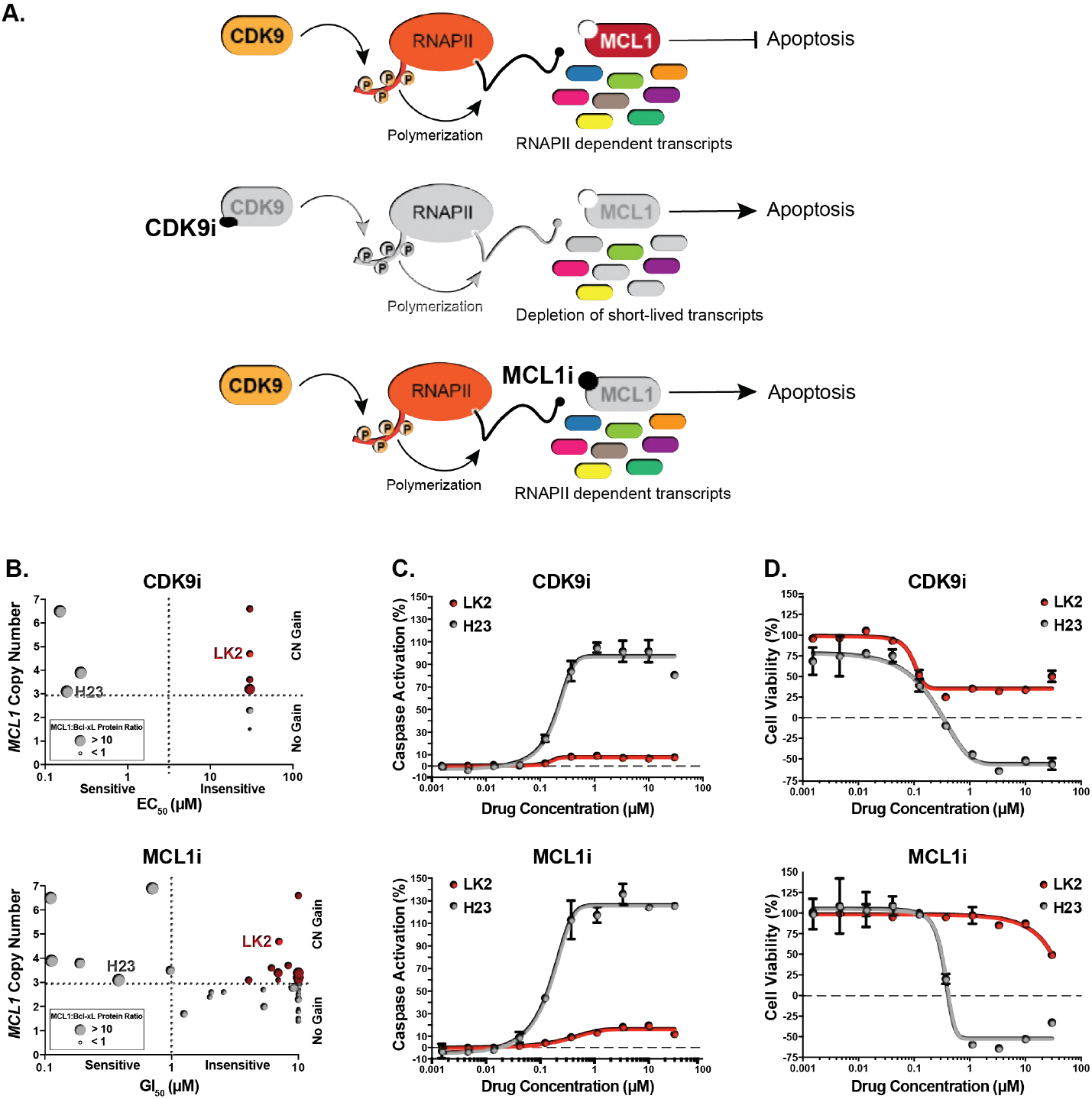
Several *MCL1*-amplified NSCLC lines are resistant to treatment with CDK9i or MCL1i. **(A)** Schematic illustrating the mechanism of action of CDK9 and MCL1 inhibitors. The CDK9 inhibitor (CDK9i) inhibits transcription elongation, thus mRNAs with short half-lives such as MCL1 are highly susceptible to acute CDK9 inhibition. The MCL1 inhibitor (MCL1i) is a BH3-mimetic that binds directly to MCL1. **(B)** Graphical representation of a panel of cell lines depicting their *MCL1* copy number, their ratio of MCL1:Bcl-xL protein and whether they are sensitive to the drug treatment indicated. EC_50_ values plotted for a 6h CDK9i treatment (top graph) derived from Caspase-Glo 3/7 assays. GI_50_ values plotted for a 24h MCL1i treatment (bottom graph) using CellTiter-Glo. Maroon circles indicate cell lines resistant to drug despite being MCL1-amplified. Highlighted in bright red is a resistant cell line (LK2) used for further study in this report and a sensitive cell line (H23) is shown in grey. **(C)** Dose response curves of LK2 and H23 treated with CDK9i (top) and MCL1i (bottom). Caspase activation was measured at 6 hours post drug treatment at the indicated concentrations by CaspaseGlo 3/7 and normalized to a positive control containing inhibitors of MCL1, BCL2 and Bcl-xL. **(D)** Cell viability curves of the resistant LK2 and sensitive H23 lines 24 hours following drug treatment with CDK9i (top) or MCL1 (bottom) at increasing concentrations as indicated. Viability was measured using the Cell Titer Glo assay normalized to a DMSO control.

Beyond direct inhibitors of the BCL2 family of proteins, inhibitors of cyclin-dependent kinase 9 (CDK9) can indirectly target MCL1. CDK9 inhibition restricts transcription elongation thus exploiting the short-lived half-life of MCL1 mRNA and protein (**Figure 1A**) (Gregory et al., 2015; C.-H. Huang et al., 2014; Lemke et al., 2014). Selective CDK9 inhibitors have been developed and show improvement in murine models of arthritis and increased survival in murine models of leukemia (Garcia-Cuellar et al., 2014; Hellvard et al., 2016). Nonetheless, these compounds must be precisely and acutely dosed for clinical applications due to their global effects on transcription. Moreover, although CDK9 inhibition suppresses MCL1 expression, it does not affect levels of some other anti-apoptotic proteins such as Bcl-xL. This exposes a potential vulnerability in treatment such that numerous cancer types may already have or develop a mode of resistance. In order to truly harness the power of CDK9 inhibitors (CDK9i) or MCL1 inhibitors (MCL1i), it is imperative to uncover additional targets that may sensitize cells to these treatments.

We reasoned that a genome wide search could uncover new targets that modulate the therapeutic activity of CDK9i and MCL1i and suggest combination therapies to resensitize tumor cells to these anti-cancer drugs. As lung cancer is the leading cause of cancer mortality and most NSCLC patients develop resistance to first-line treatment, we performed genome-wide CRISPR inhibition (CRISPRi) screens in a NSCLC line resistant to both CDK9 and MCL1 inhibition (Siegel, Miller, & Jemal, 2016). We discovered that disruption of the cullin 5-RING ubiquitin ligase (CRL5) complex markedly resensitized cells to CDK9i or MCL1i. We show that the CRL5 complex causes resistance to loss of MCL1 by targeting pro-apoptotic BH3-only proteins for proteasomal degradation. Members of the CRL5 complex are thus attractive new targets for future combination therapy to treat otherwise resistant NSCLC.

## RESULTS

### *MCL1*-amplified NSCLC lines resistant to CDK9i and MCL1i have increased Bcl-xL

We assessed a panel of NSCLC lines for their sensitivity to CDK9i (AZD5576) or MCL1i (AZD5991) currently in clinical trials (**Figure 1B and Table S1**). Several cell lines with amplified *MCL1* (copy number ≥ 3) rely on overexpression of MCL1 to escape apoptosis and were highly sensitive to CDK9i and MCL1i. We also found several NSCLC lines that remained resistant to CDK9i and MCL1i despite being *MCL1*-amplified. We examined the ratios of MCL1 protein to Bcl-xL protein and found increased Bcl-xL expression correlated with the *MCL1*-amplified lines that were resistant to CDK9i and MCL1i. Conversely, *MCL1*-amplified cell lines with low levels of Bcl-xL were sensitive to CDK9i and MCL1i.

Monitoring caspase activation in response to varying doses of MCL1i and CDK9i shows that both drugs induce apoptosis and cell death in a sensitive cell line (H23) after just six hours, with saturated cell death and caspase activation at 1μM. Conversely, a resistant line (LK2) was about 100-fold more resistant to CDK9i and MCL1i at similar drug concentrations (**Figure 1C**). Cell viability measurements illustrate the intrinsic resistance of LK2s to MCL1i with a decrease in viability observed only at the highest dose tested of 30μM, in contrast to H23s that showed a large drop in cell viability at just 0.3μM of compound. For CDK9i however, a significant decrease in cell viability is seen in both the resistant and sensitive cell lines after extended treatment (24 hours) with 0.1μM compound, highlighting the off-target toxicity of CDK9i at later time points presumably stemming from global transcriptional arrest (**Figure 1D**). This emphasizes the requirement of a tightly regulated acute treatment window for CDK9i to avoid non-specific cell death at later time points.

### Genome-wide CRISPRi screens identify factors that synergize with MCL1 inhibition

Preclinical studies suggest dual inhibition of MCL1 and Bcl-xL may not be a clinically viable option due to the highly toxic side effects. We therefore sought to identify alternate pathways that could resensitize cancer cells to CDK9i or MCL1i. LK2 cells are resistant to CDK9i and MCL1i, tolerate dense growth conditions, proliferate rapidly and transfect and transduce easily, making them an optimal cell line for a functional genomics screen. Because prolonged CDK9i treatment induces non-specific cell death, a typical growth-based screen that measures cell abundance after 14 to 21 days of compound exposure was inappropriate. Instead, we designed a positive selection FACS-based screen for apoptosis that could finish within 6 hours (the optimal treatment window to maximize on-target cell death and minimize non-specific cell death). We designed the screen to expose cells to an acute treatment of CDK9i or MCL1i and then immediately sort apoptosing cells using Cell Event staining, which fluoresces upon cellular caspase activation (**Figure 2A**).

**Figure 2.**
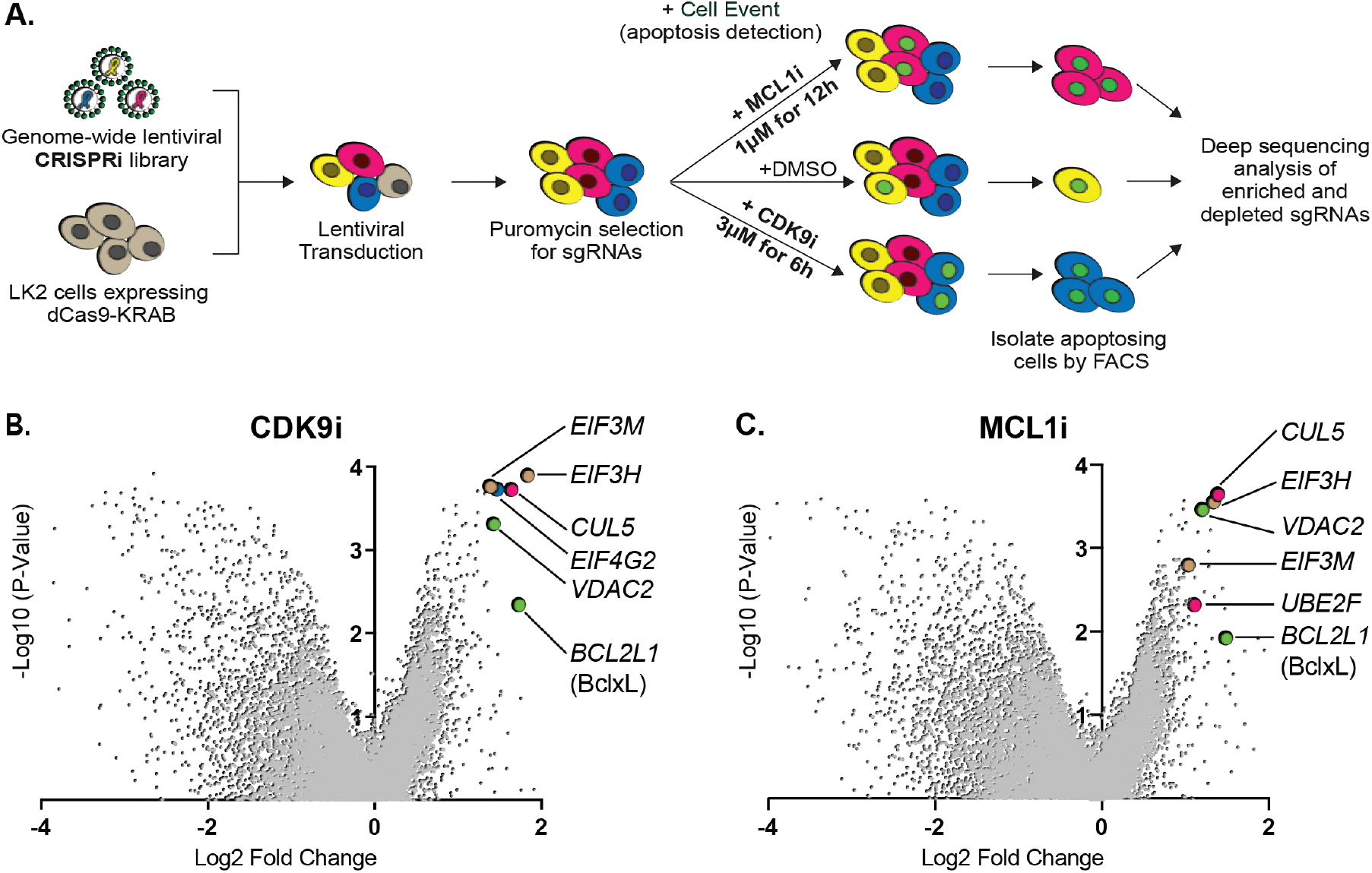
Genome-wide CRISPRi screens identify factors that resensitize lung cancer cells to inhibition of MCL1. **(A)** Schematic outlining the genome-wide CRISPRi screen in LK2 cells. Cells were exposed to acute drug treatments, fixed and FACS-sorted using the fluorogenic apoptotic detection reagent Cell Event. Enriched and depleted sgRNAs were identified by next-generation sequencing. **(B + C)** Volcano plots showing sgRNA-targeted genes significantly enriched or depleted in the apoptosing cell population following treatment with CDK9i **(B)** or MCL1i **(C)**. Average of 2 independent experiments is graphed. Green highlighted points indicate genes with a known role in apoptosis that had a significant fold change over background. Magenta points highlight members of the CUL5-RNF7-UBE2F ubiquitin complex that were significantly enriched. Beige highlighted points are members of the *E*ukaryotic *T*ranslation *I*nitiation *F*actor *3* (eIF3) complex. Blue point on CDK9i volcano plot highlights *EIF4G2*, a gene that may be involved in cap-independent translation of Bcl-xL.

We generated a clonal LK2 CRISPRi cell line to ensure uniform expression of dCas9-KRAB and functionally validated knockdown of a panel of genes by qRT-PCR (**Figure S1A**). Biological duplicates of LK2 CRISPRi cells were transduced with a genome-wide sgRNA library at a low multiplicity of infection to ensure delivery of one sgRNA per cell, and at 500X coverage to promote equal guide representation. The sgRNA vectors also encode a puromycin resistance cassette allowing for puromycin-selection of successfully transduced cells. Library transduced cells were treated with either 3μM CDK9i for 6 hours or 1μM MCL1i for 12 hours and incubated with 0.5μM Cell Event for the duration of the treatment. The concentrations and length of treatments were determined based on where maximal differences in caspase activation were observed between resistant and sensitive cell lines. To control for vehicle effects, we also performed two duplicate genome-wide sorts with DMSO as a treatment condition. We harvested bulk untreated cells on the same day as the FACS sort of DMSO- or drug-treated cells to provide an accurate ‘background’ sampling of sgRNAs and eliminate confounding effects from essential genes that had dropped out of the population. We measured quantitative differences in sgRNA frequency by deep sequencing and integrated enriched or depleted sgRNAs into gene-level hits by comparing sorted samples to the untreated control using ScreenProcessing (**Table S2-S4**) and MAGeCK (**Table S5-S8**) analysis pipelines (Horlbeck et al., 2016; Li et al., 2014).

We analyzed DMSO-treated apoptosing cells to determine if any gene knockdowns promoted apoptosis upon exposure to vehicle. No significant hits in the DMSO control were detected over the random distribution of non-targeting sgRNA controls (**Figure S1B**), indicating that our gene calls in CDK9i- or MCL1i-treated samples were specifically derived from drug synergy. We found multiple sgRNAs targeting Bcl-xL led to re-sensitization of LK2 cells to CDK9i and MCL1i (**Figure 2B-C**), confirming reports that depletion of both MCL1 and Bcl-xL initiates apoptosis (Goodwin et al., 2015; Xiao et al., 2015; Zhang et al., 2011). Downregulation of the mitochondrial porin VDAC2 also promoted apoptosis in drug-treated cells (**Figure 2B-C**), consistent with its proposed role in BAK/BAX sequestration at the mitochondrial membrane (Cheng, Sheiko, Fisher, Craigen, & Korsmeyer, 2003; Lauterwasser et al., 2016) (Chin et al., 2018). Interestingly, *EIF4G2* (DAP5) was identified as a hit in the CDK9i screen (**Figure 2B**) and has been implicated in stimulating cap-independent translation of anti-apoptotic protein BCL-2, potentially skewing the balance in favor of apoptosis (Marash et al., 2008).

While CDK9i and MCL1i reduce MCL1 activity through completely separate mechanisms, the top resensitization hits were strikingly consistent between both screens. These hits are furthermore part of two physical complexes, one potentially involved in specialized translation and the other involved in protein degradation. First, knockdown of multiple components of the eukaryotic translation initiation factor 3 (eIF-3) complex (EIF3H and EIF3M) resensitize LK2 cells to CDK9i and MCL1i (**Figure 2B-C and Figure S1C-D**). eIF-3 is reported to bind a highly specific program of messenger RNAs involved in cell proliferation and apoptosis (Lee, Kranzusch, & Cate, 2015). One might speculate that this complex could be involved in mediating translational activation of certain anti-apoptotic proteins or repression of pro-apoptotic proteins. Regulation of apoptosis by eIF-3 warrants further investigation, but we opted to focus on the second, better-studied complex.

Knockdown of multiple members of a cullin-RING ubiquitin ligase complex (CRL) including CUL5, UBE2F and RNF7 resensitizes LK2 cells to CDK9i and MCL1i (**Figure 2B-C and Figure S1C-D**). The cullin 5 (CUL5) scaffold was identified as a top resensitizing hit in both screens. Repression of the NEDD8-conjugating activation subunit UBE2F sensitizes LK2 cells to MCL1i (**Figure 2C and Figure S1D**). Additionally, knockdown of Ring Finger Protein 7 (RNF7), a protein that binds to the CUL5 scaffold and serves as a catalytic subunit, sensitizes cells to both CDK9i and MCL1i (**Figure S1C-D**).

### Depletion of the CUL5-RNF7-UBE2F ubiquitin ligase complex resensitizes cancer cells to treatment with CDK9i or MCL1i

Cullin-RING ligases are emerging as attractive cancer targets and a novel class of small molecule neddylation inhibitors have recently been developed (Soucy et al., 2009). While the cullin 3 complex (CRL3) is more typically associated with cancer phenotypes, there is little data on a role for CRL5 in tumorigenesis or resistance. We therefore sought to further examine the role of the CRL5 complex in resensitizing LK2 cells to CDK9 and MCL1 inhibition.

Cullin-RING ubiquitin ligases (CRLs) comprise the largest known category of ubiquitin ligases and are involved in regulating numerous dynamic cellular processes (Petroski & Deshaies, 2005). They are multi-subunit complexes built upon the cullin scaffold proteins, and CUL5 plays this role in the CRL5 complex (**Figure 3A**). CUL5 binds to a RING protein (RNF7) that serves as a docking site for E2 ubiquitin ligases. CUL5 binds exclusively to RNF7, indicating that RNF7 is indispensable for CRL5 activity (Kamura et al., 2004). UBE2F, also known as the NEDD8-conjugating enzyme (NCE2) is an E2 ubiquitin ligase that neddylates and activates CRL5 (D. T. Huang et al., 2009). Elongin B and C alongside a substrate adaptor bind to the N-terminus of CUL5 and dictate choice of substrates that get polyubiquitinated by E3 ubiquitin ligases (Mahrour et al., 2008; Okumura, Joo-Okumura, Nakatsukasa, & Kamura, 2016). Adaptor proteins add a layer of regulation to CRL activity both by specifying substrates as well as defining the timing and localization of ubiquitination, however the substrate adaptor regulating CRL5 resistance to MCL1 inhibition is not known.

**Figure 3.**
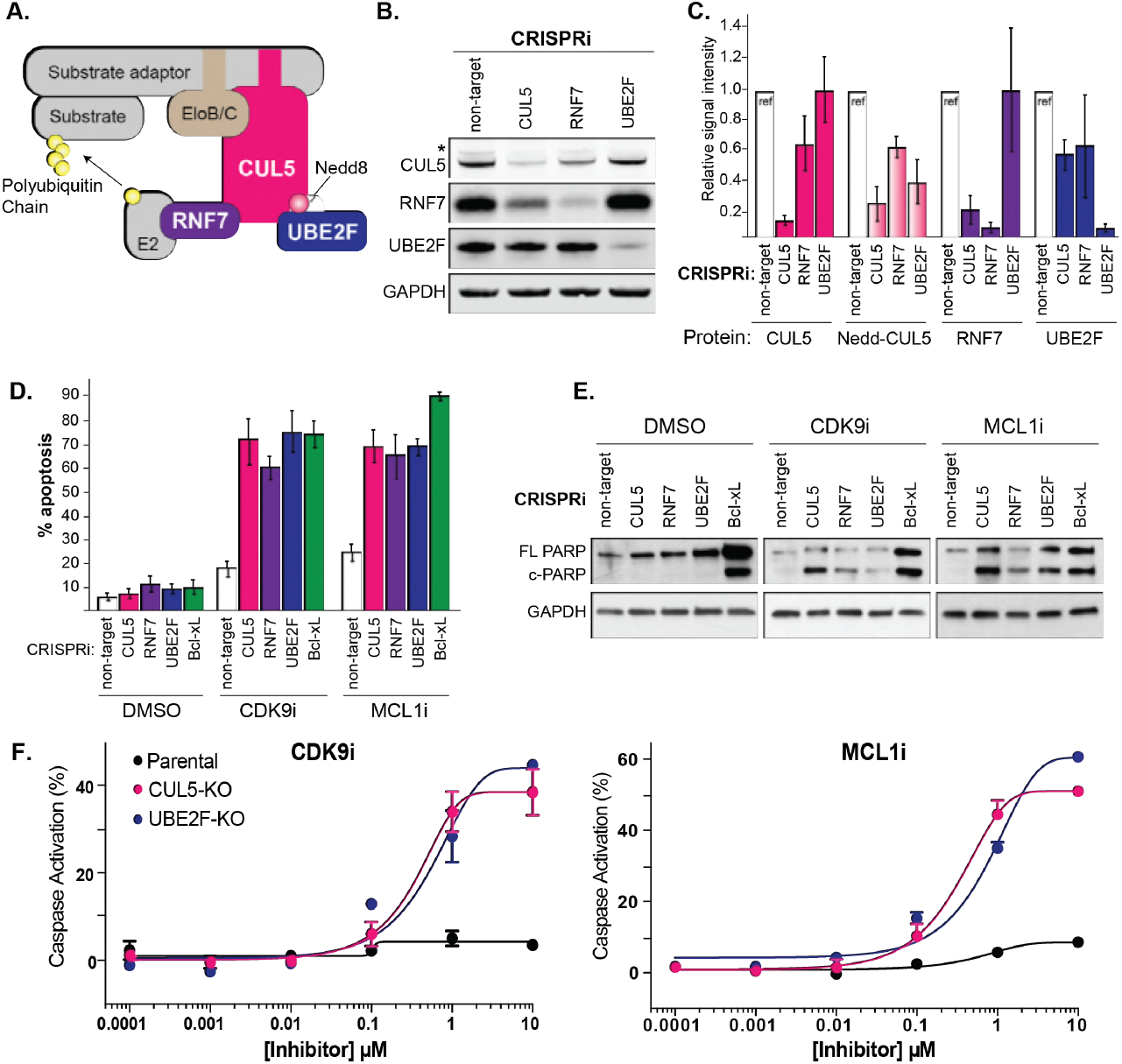
Depletion of the CUL5-RNF7-UBE2F ubiquitin complex induces apoptosis upon treatment with CDK9i or MCL1i. **(A)** Schematic depicting the CUL5 ubiquitin ligase scaffold and its interacting partners. **(B)** Western blots confirming effective knockdown of CUL5, RNF7 and UBE2F by stable lentiviral expression of dCas9-KRAB and corresponding sgRNAs. Asterisk indicates neddylated CUL5. GAPDH serves as loading control. **(C)** Quantification of western blots as in B. Error bars show standard deviations from three independent biological replicates. **(D)** Induction of apoptosis when cells depleted of CUL5, RNF7 or UBE2F (as in B) are treated with CDK9i (3μM for 6h), MCL1i (1μM for 12h) or DMSO. Knockdown of Bcl-xL serves as a positive control for induction of apoptosis. Percentage of apoptosis determined by flow cytometry detection of the fluorogenic apoptotic detection reagent, Cell Event. Error bars are standard deviations of three independent biological replicates. **(E)** Western blotting for cleaved PARP (c-PARP) serves as orthogonal readout for induction of apoptosis following knockdown of target genes and treatment with 3μM CDK9i for 6 hours or 1μM MCL1i for 12 hours. Knockdown of Bcl-xL is included as a positive control, however due to extreme toxicity of the combination treatment, cells were harvested 3 hours after treatment with 3μM CDK9i or 0.5 hours after treatment with 1μM MCL1i. GAPDH, loading control. **(F)** Dose response curves of caspase induction showing resensitization of CUL5-knockout (CUL5-KO c3) and UBE2F-knockout (UBE2F-KO c1) lines as compared to parental LK2 cells when treated with CDK9i (top) and MCL1i (bottom). Caspase activation was measured at 10 hours post drug treatment at the indicated concentrations by CaspaseGlo 3/7 and normalized to a positive control containing inhibitors of MCL1, BCL2 and Bcl-xL.

Interestingly, another hit in the MCL1i screen was the BTB/POZ domain-containing protein 3 (BTBD3; **Figure S1D**). While BTBD3 has no known function, members of the BTB (also known as POZ) domain containing protein family can act as substrate adaptors for cullin 3 (L. Xu et al., 2003). We speculated that BTBD3 might be a substrate adaptor for CUL5 that specifically mediates resensitization to MCL1i and CDK9i, and thus included it in later validation.

We made stable dCas9-KRAB LK2 cell lines that expressed an individual CRISPRi sgRNA to knockdown CUL5, RNF7, UBE2F, as well as a non-targeting (NT) control. We confirmed efficient knockdown of CUL5, RNF7 and UBE2F by western blotting (**Figure 3B**). CUL5 and RNF7 appeared to be dependent on each other for stability such that when either protein was knocked down, levels of the corresponding protein were also diminished (**Figure 3C**). Depletion of UBE2F resulted in the disappearance of a higher molecular weight band corresponding to neddylated CUL5 (**Figure 3B-C**).

Using two independent stable LK2 CRISPRi lines targeting CUL5, RNF7, UBE2F or Bcl-xL, we validated that individual knockdowns of these genes followed by treatment with CDK9i or MCL1i induced extensive caspase activation (**Figure 3D and Figure S2A**). Bcl-xL served as a positive control based on previous reports that dual inhibition of Bcl-xL and MCL1 induces apoptosis (Goodwin et al., 2015; Xiao et al., 2015; Zhang et al., 2011). As an orthogonal readout for apoptosis, we also assessed cleaved PARP (c-PARP) levels following drug treatment (**Figure 3E**). No c-PARP was observed in untreated stable knockdowns of CUL5, RNF7 or UBE2F, indicating that the CRL5 complex is not intrinsically required for LK2 cell survival. By contrast, knockdown of Bcl-xL alone led to substantial apoptosis (**Figure 3E**). Depletion of CUL5, RNF7, UBE2F or Bcl-xL when combined with CDK9i or MCL1i significantly induced PARP cleavage (**Figure 3E**). Hence, loss of CRL5 activity alone does not induce apoptosis, suggesting that it could be less toxic than Bcl-xL inhibition and better suited to co-administration with MCL1 inhibitors.

To further validate the synergy observed between inhibition of CRL5 and MCL1, we made isogenic knockouts of CUL5 and UBE2F. Purified Cas9 ribonucleoproteins (RNPs) complexed with sgRNA pairs flanking coding exon 1 for CUL5 or UBE2F were nucleofected into LK2s in order to completely disrupt coding potential (Lingeman, Jeans, & Corn, 2017). Nucleofected pools were seeded at single cell densities by FACS sorting and clones were initially screened for loss of protein by western blot and confirmed by Sanger sequencing (**Figure S2B-C**). We did not attempt to knockout RNF7 because cells did not tolerate extended culturing of CRISPRi-mediated stable knockdowns of RNF7. Knockouts of CUL5 and UBE2F were viable and displayed high levels of apoptosis when challenged with a single concentration of CDK9i or MCL1i (**Figure S2D**). Dose response curves in the CUL5 and UBE2F knockouts showed significant caspase activity at 1μM of CDK9i or MCL1i, whereas wild type cells showed no response even at 10μM doses (**Figure 3F**). Taken together, multiple lines of evidence indicate that depletion of the CRL5 complex does not on its own induce high levels of apoptosis, but resensitizes cells to CDK9 and MCL1 inhibition.

To further establish the mechanistic basis for CRL5 protection from CDK9i and MCL1i, we asked whether depletion of Elongin B, Elongin C, or the potential BTBD3 adapter also synergized with MCL1 inhibition. We performed individual knockdowns of each gene by CRISPRi (**Figure S3A**). Reduction of Elongin B but not Elongin C induced apoptosis at levels similar to that of CUL5 knockdown (**Figure S3B**). Elongin B and C have often been described as functioning in a complex (Okumura, Matsuzaki, Nakatsukasa, & Kamura, 2012), but our results suggest a potential separation of function of the two proteins in sensitivity to MCL1 inhibitors. Knockdown of BTBD3 showed a modest increase in apoptosis when cells were treated with MCL1i but did not entirely phenocopy CUL5 or Elongin B knockdown (**Figure S3B**). This suggests that even if BTBD3 does play a role in the CRL5 complex, it does not appear to be the primary substrate adaptor directing ubiquitination of a substrate(s) that synergizes with MCL1 inhibition to induce apoptosis.

In an attempt to find the CRL5 substrate adaptor responsible for resensitization to MCL1 inhibitors, we used CRISPRi to individually knock down a large number of CUL5 substrate adaptors annotated in literature and tested them for synergy with MCL1i and CDK9i (Okumura et al., 2016). Two different sgRNAs were tested per gene, but none of the putative substrate adaptors analyzed induced apoptosis following CDK9i or MCL1i treatment (**Figure S3C and Table S9**). We cannot rule out redundancy between substrate adaptors such that one compensates for loss of another. The relevant CRL5 substrate adaptors in the context of resensitization to MCL1 inhibition remain to be identified.

### The CRL5 Complex Regulates Levels of the BH3-only Apoptotic Sensitizers Bim and Noxa

We next searched for the substrate of CRL5 that potentiates resensitization to MCL1 inhibition. A literature search for apoptosis-related substrates of CUL5 implicated p53, a master regulator of apoptosis. Several viral proteins hijack the CUL5 complex and serve as substrate adaptors that bind p53 and mediate its degradation, which promotes viral infection (Cai, Knight, Verma, Zald, & Robertson, 2006; Querido et al., 2001; Sato et al., 2009). If the CRL5 complex degrades p53 in LK2 cells, inactivating CRL5 would result in accumulation of p53 thereby promoting apoptosis. We Western blotted for p53 in drug treated wild type and CUL5-knockdown cells but observed no difference in p53 levels (**Figure S3D**). We also confirmed by western blotting that depletion of CUL5 had no effect on MCL1 and Bcl-xL levels (**Figure S3E**).

Since p53 levels were not directly affected by loss of CRL5, we hypothesized that CRL5 could be targeting pro-apoptotic proteins for proteasomal degradation. In this case, loss of CRL5 would result in accumulation of pro-apoptotic factors. This accumulation alone is presumably not sufficient to induce apoptosis due to the overexpression of anti-apoptotic proteins MCL1 and Bcl-xL. However, when combined with inhibition of MCL1, the balance may then be skewed in favor of apoptosis. Indeed, Noxa was recently proposed to be a substrate of CRL5, but this molecular activity was not linked to a cellular phenotype (Zhou et al., 2017).

We used Western blotting to interrogate the levels of all eight BH3-only proteins in individual CRISPRi knockdowns of CUL5, RNF7, UBE2F and a NT control. Strikingly, we observed increased protein levels of only two BH3-only proteins, Noxa and Bim (**Figure 4A**). To determine whether treatment with CDK9i or MCL1i revealed any additional differences in protein levels, we treated NT or CUL5 knockdown cells with DMSO, CDK9i or MCL1i and immunoblotted for all BH3-only proteins. We found that levels of both proapoptotic proteins Noxa and Bim were higher in cells lacking CUL5, while all other BH3-only proteins remained unchanged (**Figure 4B-C**). Interestingly, CDK9i alone reduced levels of Noxa and Bim, and this occurred through transcriptional downregulation that is presumably related to the half-lives of these mRNAs (**Figure 4D**). Knockdown of CUL5 together with CDK9i rescued protein levels of Noxa and Bim but not mRNA levels, consistent with a model in which CRL5-mediated post-translational degradation of Noxa and Bim is the mechanism of resistance to MCL1 inhibition.

**Figure 4.**
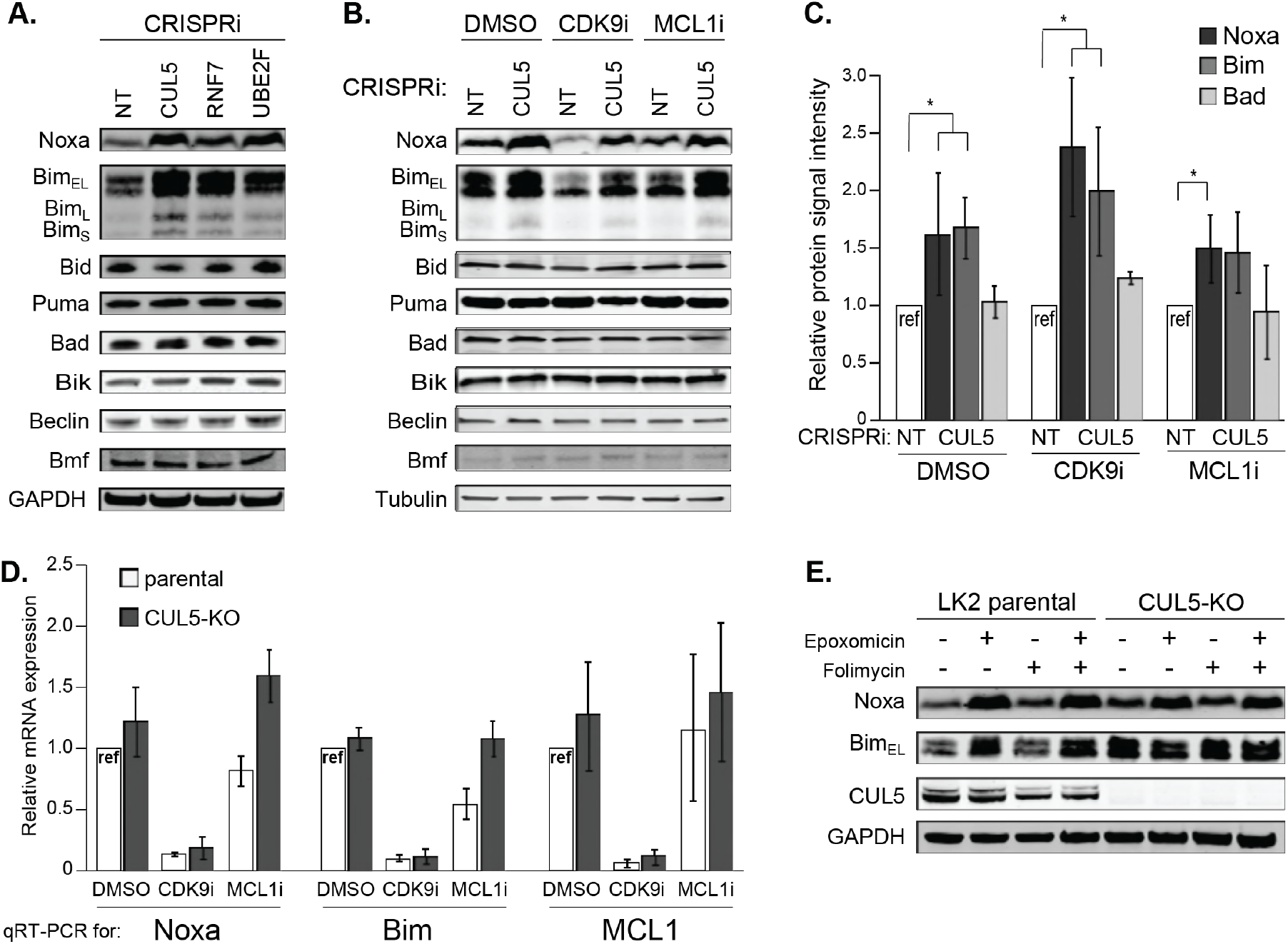
The CUL5-RNF7-UBE2F ubiquitin complex regulates levels of BH3-only apoptotic sensitizers Bim and Noxa. **(A)** Western blotting of all BH3-only proteins in cell lines with depleted CUL5, RNF7 or UBE2F show increased levels of Bim and Noxa as compared to cells transduced with a non-targeting (NT) sgRNA. GAPDH serves as the loading control. **(B)** Western blot of all BH3-only proteins in NT and CUL5-depleted cells treated with DMSO, CDK9i (3μM for 6h) or MCL1i (1μM for 12h). Tubulin serves as loading control. **(C)** Quantification of western blots as in (B). Error bars derived from standard deviations of three biological replicates consisting of independently treated and isolated protein samples. Asterisk indicates p value <0.05 as determined by a paired student T-Test. **(D)** qRT-PCR showing relative mRNA levels of Noxa, Bim and MCL1 in LK2 parental and CUL5-KO c1 cells treated with DMSO, CDK9i (3μM for 6h) or MCL1i (3μM for 6h). Error bars show standard deviations from three biological replicates on independently treated and isolated RNA samples. **(E)** Western blot of LK2 and CUL5-KO c1 cells treated for 6 hours with epoxomicin (100μM), or folimycin (100μM) or both epoxomicin and folimycin (100μM each). GAPDH, loading control.

To confirm that CRL5 targets Noxa and Bim for proteasomal degradation, we inhibited proteaosmal and lysosomal function with epoxomicin and folimycin, respectively. No increases in Bim or Noxa were detected following folimycin treatment, indicating that these proteins are not lysosomally degraded (**Figure 4E**). We found that Bim accumulates in epoxomicin-treated LK2 parental cells at levels equivalent to untreated CUL5-KO cells. No change in Bim accumulation was observed in CUL5-KO cells treated with epoxomicin. These results suggest that Bim is targeted for proteasomal degradation exclusively by CUL5 in LK2 cells, as Bim levels are completely rescued by inhibition of the proteasome in wild type cells but proteasomal inhibition has no effect on Bim levels in CUL5-KO cells. Noxa accumulates in both parental and CUL5-KO cells following epoxomicin treatment (**Figure 4E**), demonstrating that Noxa is also proteasomally regulated. However, inhibition of the proteasome also rescues Noxa levels in CUL5-KO cells, suggesting that Noxa can be targeted by additional ubiquitin ligase complexes. We were unable to demonstrate direct targeting of Noxa and Bim by CRL5 using ubiquitin co-immunoprecipitation (**Figure S4**), leaving open the possibility that these proteins are indirect substrates of the CRL5 ligase complex.

### Noxa is Required to Resensitize CUL5-deficient cells to MCL1 Inhibition

Based on the observation that removal of CUL5 alone does not induce apoptosis despite increased levels of pro-apoptotic Bim and Noxa, we reasoned the high expression of MCL1 in these cells was still sufficient to prevent cell death. However, when MCL1 is inhibited in cells lacking CUL5, the accumulated Bim and Noxa may now be able to initiate apoptosis.

To examine the dependency of Noxa and Bim to resensitize CUL5-deficient cells to CDK9i or MCL1i, we used RNAi to knock down Noxa and/or Bim in the context of CUL5 CRISPRi. If induction of apoptosis in CUL5-deficient cells requires Noxa and/or Bim, then removal of either or both factors should limit apoptosis and make the cells re-resistant to inhibition of MCL1. To validate the experimental design, we co-depleted CUL5 and BAK, the apoptosis effector required for cell death. In these conditions, treatment with CDK9i or MCL1i no longer induced apoptosis, consistent with the requirement of BAK in the initiation of apoptosis (**Figure 5A-B and Figure S5A-B**). Induction of apoptosis remained high in MCL1i-treated cells lacking Noxa and CUL5 or cells lacking Bim and CUL5. Similarly co-depletion of all three proteins Noxa, Bim and CUL5 did not restrict apoptosis upon MCL1i treatment (**Figure 5B and Figure S5B**). Thus Bim and Noxa are insufficient to induce cell death when CUL5-depleted cells are treated with MCL1i.

**Figure 5.**
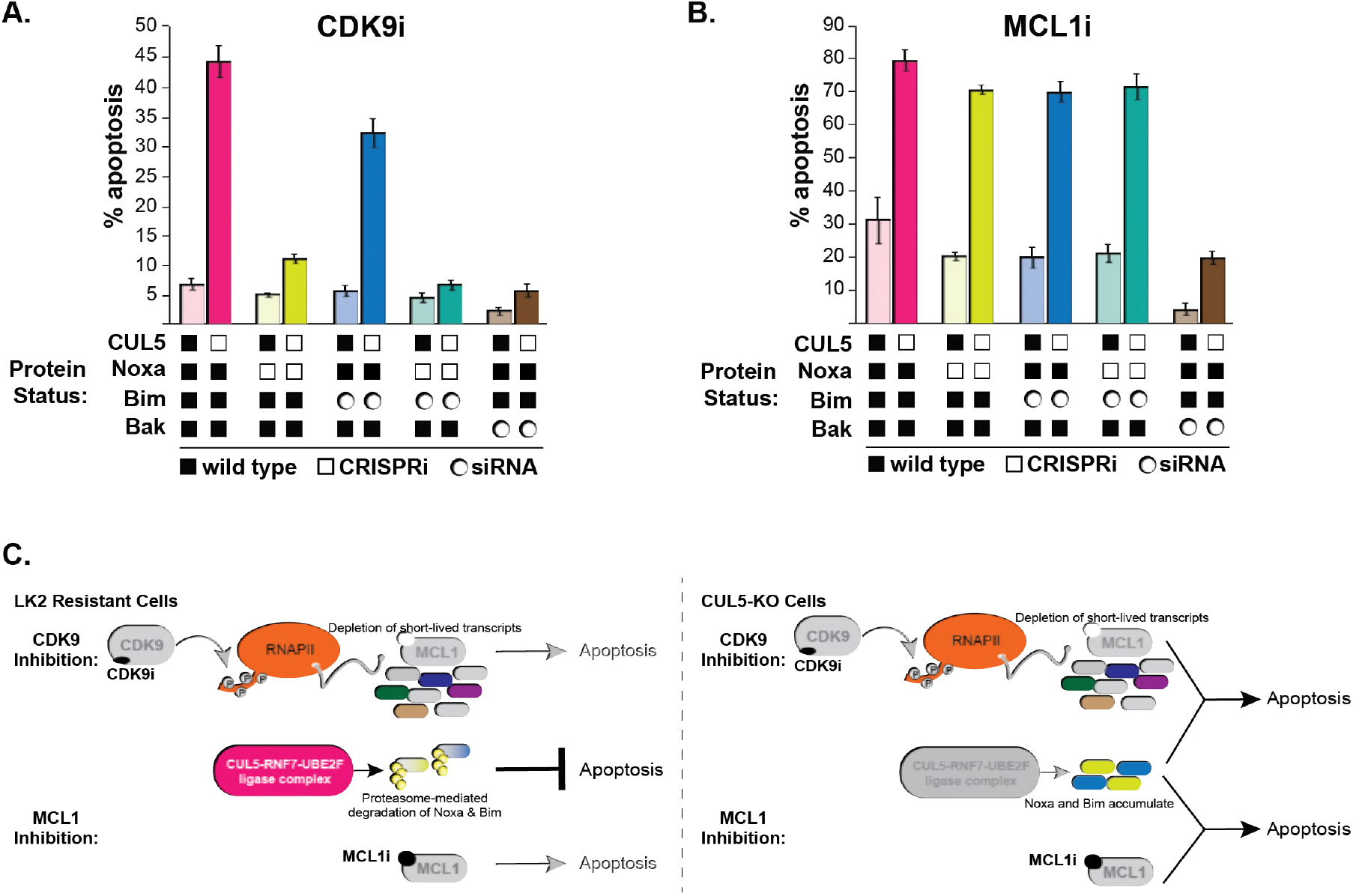
CUL5-depleted cells regain resistance to CDK9i, but not MCL1i, when Noxa is knocked down. **(A)** CUL5 and Noxa were knocked down in dCas9-KRAB-expressing LK2 cells by selecting for corresponding sgRNAs. Bim and Bak were knocked down by siRNA. Apoptosis was measured using Cell Event detection by flow cytometry after treatment with 3μM CDK9i for 6 hours. Error bars are standard deviations of 3 biological drug treatment replicates. **(B)** Genetic manipulations were performed and apoptosis was measured as in (A) following MCL1i treatment (1μM for 12 hours). **(C)** Schematic illustrating potential mechanisms of resensitization of resistant NSCLC lines to inhibition of MCL1.

However, CDK9i treatment of Noxa and CUL5 co-depleted cells led to almost no apoptotic induction, indicating these cells had regained resistance to CDK9i (**Figure 5A and Figure S5A**). Removal of both Noxa and Bim in CUL5-depleted cells did not lead to an additive effect in the reacquisition of resistance to CDK9i, while cells lacking only Bim and CUL5 were still sensitive to CDK9i. These data indicate that in CDK9i-treated cells lacking CUL5, initiation of apoptosis is primarily dependent on Noxa.

We wondered whether the discrepancy between MCL1i and CDK9i treatments was caused by the additional depletion of Noxa and Bim due to non-specific effects of CDK9i that we previously observed (**Figure 4B-C**). However qRT-PCR after knockdown of Noxa and Bim already showed extremely low levels of Noxa and Bim mRNA (**Figure S5C**), making this an unlikely explanation. Instead, the difference in MCL1i and CDK9i may arise from CDK9i targeting an additional protein involved in promoting apoptosis, leading to overall abrogation of apoptosis. In conclusion, our work reveals that resistance to CDK9i or MCL1i can be overcome by inactivation of the CRL5 complex and we propose this is in part mediated by the accumulation of the CRL5 target Noxa (**Figure 5C**).

## DISCUSSION

Cullin-RING complexes comprise the largest class of ubiquitin ligases and regulate a diverse array of biological processes. While CUL3 is implicated in a wide array of cancer biologies, relatively little is known about the physiological functions of many CUL5-containing ubiquitin ligases. A few reports have implicated CUL5 in cancer progression. Downregulation of CUL5 has been observed in numerous breast tumors (Fay et al., 2003) and overexpression of CUL5 in T47D breast cancer cells has been shown to limit cell growth (Burnatowska-Hledin et al., 2004; Lubbers et al., 2011). Additionally, microRNAs 19a/b that negatively regulate CUL5 promote cell growth and invasion in cervical cancer, which is abolished by complementation with miRNA-resistant CUL5 (X.-M. Xu et al., 2012). While alterations in CUL5 are correlated with cancer phenotypes, no clear mechanism has emerged to explain these findings. Our work reveals the CUL5-RNF7-UBE2F ubiquitin ligase as a key modulator of apoptosis and suggests that CUL5 may restrict apoptosis by the constitutive degradation of Noxa and/or Bim. Indeed, previous work has shown that the CUL5-RNF-UBE2F complex is capable of *in vitro* ubiquitination of Noxa (Zhou et al., 2017). While several viral proteins hijack the CUL5 complex to degrade p53, a master regulator of apoptosis initiation (Cai et al., 2006; Querido et al., 2001; Sato et al., 2009), we found no evidence however of differential p53 levels in DMSO-, CDK9i- or MCL1i-treated wild type and CUL5-KO cells. Thus, we suggest that p53 signaling is not a major mechanism involved in resensitization of cells to MCL1 inhibition.

Our discovery that inactivation of the CRL5 complex can resensitize cancers to MCL1 inhibition provides a therapeutic opportunity, not only in the identification of novel drug targets, but also as a potential biomarker for patient stratification. Based on our data, lung cancer patients that acquire loss of function mutations in either CUL5, RNF7, UBE2F or Elongin B that coincide with high expression of MCL1/Bcl-xL would be candidates for treatment with CDK9i or MCL1i because they are unlikely to have pre-existing CRL5-mediated resistance or to acquire such resistance during treatment. In fact we note that there is significant co-occurrence of MCL1 amplifications with CUL5 mutations and UBE2F deletions in NSCLC adenocarcinomas deposited in The Cancer Genome Atlas (TCGA) (Cerami et al., 2012; Gao et al., 2013). Additionally, as MCL1 and Bcl-xL focal amplifications co-occur frequently in breast cancer, it is possible that the same stratification strategy may apply to a subset of breast cancer patients as well to NSCLC.

Currently no small-molecule inhibitors exist to target specific components of the CRL5 complex. However, several potent inhibitors that broadly inhibit ubiquitin-proteasome systems have been developed. MLN4924 inhibits the NEDD8 activating enzyme (NAE), thereby preventing conjugation of NEDD8 onto all cullin scaffolds, and shows significant anti-tumor activity in clinical trials (Soucy et al., 2009). Alternatively, there are several clinically approved proteasome inhibitors such as bortezomib and carfilzomib currently used in the treatment of multiple myeloma and lymphomas (Richardson et al., 2005) (Kuhn et al., 2007). Although a therapeutic window exists for these compounds, patients also experience adverse side effects due to the widespread effects of proteasome inhibition, making it a complicated option for co-dosing strategies with CDK9i or MCL1i.

Recently more attention has focused on the development of compounds that directly disrupt protein-protein interactions within specific Cullin-RING ligases, with promising results. One example is the generation of a potent small molecule targeting the von Hippel-Landau (VHL) substrate adaptor (Galdeano et al., 2014). VHL functions in complex with CRL2 to degrade an essential transcription factor HIF-1α, which promotes erythropoietin production. Other broad-spectrum drugs that prevent HIF-1α degradation are in late-stage clinical trials for treatment of chronic anemia associated with kidney disease and chemotherapy (Joharapurkar, Pandya, Patel, Desai, & Jain, 2018). Due to the specialized composition of individual CRL complexes associated with distinct biological functions, this approach promises a more specific and hopefully less toxic inhibitor. Moreover, our observation that CUL5 and UBE2F knockouts are viable suggests inhibition of these components may be well tolerated making them ideal co-dosing targets. Identification of the substrate adaptor responsible for CRL5-mediated regulation of Noxa/Bim could yield a promising target for MCL1-addicted tumors.

Innate and acquired resistance to MCL1 inhibition remains a major challenge for lung (and breast) cancer patients, often due to the concomitant amplification of additional anti-apoptotic factors such as Bcl-xL. Targeting multiple anti-apoptotic proteins may limit the therapeutic window due to the accompanying cytotoxicity. Our results reveal an alternate pathway that may be targeted in combination with MCL1 inhibition to combat resistance and hopefully provide a safe and effective therapy.

## Author Contributions

SK and JEC designed experiments. JC, CA, CD and MB assessed panel of cell lines for MCL1/Bcl-xL status and sensitivity to CDK9i and MCL1i; they also performed all dose-response and viability curves. PCL constructed and validated the initial CRISPRi clonal LK2 cell line. SK performed all the screens and follow-up experiments. TM analyzed screen results with input from SK, JC, JEC and LD. HM assisted with the validation experiments. RB generated the CUL5 and UBE2F knockout clones. SK and JEC wrote the manuscript with input from JC.

## Funding Acknowledgements

This work was supported by a National Institutes of Health New Innovator Awards (DP2 HL141006), the Li Ka Shing Foundation, the Heritage Medical Research Institute, the California Institute of Regenerative Medicine (DISC1-08776), and funded research support from AstraZeneca. This work used the Vincent J. Coates Genomics Sequencing Laboratory at UC Berkeley, supported by the NIH S10 OD018174 Instrumentation Grant.

## MATERIALS AND METHODS

### Cell culture and reagents

The LK2 cell line was obtained from AstraZeneca and maintained in RPMI supplemented with 10% (v/v) fetal bovine serum, 100 units/mL penicillin and streptomycin, GlutaMAX, sodium pyruvate and non-essential amino acids. All cells tested negative for Mycoplasma contamination using the MycoAlert Plus Mycoplasma detection kit from Lonza. AstraZeneca licensed the CDK9 inhibitor (AZD5576) originally published as PC585 (Garcia-Cuellar et al., 2014) and supplied it to us for this study. The MCL1 inhibitor (AZD5991) is synthesized by AstraZeneca (Tron et al., 2018).

### Lentivirus production and transduction

Lentivirus was produced by transfecting HEK293T with standard packaging vectors using the *Trans*IT-LT1 Transfection Reagent (MIR 2306; Mirus Bio LLC). Viral supernatant was collected at 48h and 72 h after transfection, filtered through a 0.45-μm polyethersulfone syringe filter, snap-frozen and stored at −80 °C for future use. For screens, viral titrations were performed by transducing LK2 cells at serial dilutions and assessing BFP (present on the sgRNA-encoding plasmid) percentages 48h following transduction. The viral dilution resulting in ~15% BFP-positive cells was used to transduce cells for the screens.

### Individual analysis of sgRNA phenotypes

sgRNA protospacers targeting CUL5, RNF7, UBE2F, Elongin B, Elongin C, Bcl-xL, Noxa and a negative control non-targeting protospacer were individually cloned into BstXI/BlpI-digested pCRISPRia-v2 (Addgene #8432) by ligating annealed complementary synthetic oligonucleotide pairs. The sgRNA expression vectors were packaged into lentivirus as previously described and successful transductants were selected with puromycin at a final concentration of 2.5μg/mL. Sequences for protospacers are in Table S10.

### Genome-scale CRISPRi screens

The human CRISPRi v2 library contains 5sgRNAs/annotated TSS of each gene comprising a total of 104,535 sgRNAs. In order to maintain 500X coverage (representation) of each guide at all times, ~50×10^6^ cells need to be transduced with one sgRNA. To ensure delivery of one sgRNA/cell a low MOI of ~0.3 is used, which results in not every cell getting transduced, thus 5-6 fold more cells need to be transduced in order to maintain 500X coverage. Replicate cultures of 3×10^8^ cells were plated in twenty 15cm dishes and transduced at an MOI of ~0.3 in the presence of 4μg/mL polybrene. Cells were split into 2.5μg/mL puromycin and selected for 4 days. Cells were passaged into regular media and recovered for 2 days. 60×10^6^ were collected as the ‘background’ sample. 60×10^6^ cells were treated with 3μM CDK9i and 0.5μM Cell Event for 6 hours. 60×10^6^ cells were treated with 1μM MCL1i and 0.5μM Cell Event for 12 hours. 60×10^6^ cells were treated with DMSO (1:10,000X dilution) and 0.5μM Cell Event for 12 hours. After treatment, cells were trypsinized, washed and fixed in 1% formaldehyde/PBS. The entirety of the cell population was then FACS-sorted where GFP-positive (apoptosing) cells were isolated. As previously mentioned, this was done in duplicate. Genomic DNA was purified from each cell population with blood purification kits (Machery-Nagel, Nucleospin blood L or XL, depending on cell number) and the sgRNA-encoding region was enriched, amplified, and processed for sequencing on the Illumina Hiseq 2500. TruSeq index sequences unique to each cell population were used to multiplex samples.

### Pooled screen analysis

Data analysis was performed as described (Horlbeck et al., 2016; Li et al., 2014). Briefly, sequence reads from the Illumina HiSeq 2500 were trimmed, aligned to CRISPRi v2 sgRNA library, counted and normalized. For the Python-based ScreenProcessing pipeline, sgRNA phenotypes and negative control gene phenotypes were determined along with Mann-Whitney P values. The top three guide-level phenotypes were collapsed to produce the gene-level phenotype score. For genes with multiple annotated transcription start sites (TSSs), phenotypes were calculated for each TSS and the TSS with the lowest Mann Whitney p-value was used to represent the gene. For MAGeCK, all sgRNAs/gene (including both TSSs if there are two) are ranked based on p-values from mean variance modeling. A robust ranking aggregation (RRA) algorithm is then used to call genes that are significantly enriched or depleted based on p-value and false discovery rate (FDR).

### qRT-PCR

For qPCR, RNA was extracted with the RNeasy Mini Kits (Qiagen). cDNA was produced from 1 μg of purified RNA using the iScript Reverse Transcription Supermix for RT-qPCR (Bio-Rad Laboratories). qPCR reactions were performed with the SsoAdvanced Universal SYBR Green Supermix (Bio-Rad Laboratories) in a total volume of 10 μl with primers at final concentrations of 500 nM. Primer sequences are included in Table S11. The thermocycler was set for 1 cycle of 95 °C for 30 sec, and 40 cycles of 95 °C for 5 s and 55 °C for 15 s, respectively. Fold enrichment of the assayed genes over the control *HPRT* and/or *GAPDH* loci were calculated using the 2^−ΔΔ*C*^T method as previously described (Livak & Schmittgen, 2001).

### Cell Event apoptosis detection assay

Cell Event reagent was added to cells for the duration of the drug or vehicle treatment, at a final concentration of 0.5μM. Cells were harvested by trypsinization and analyzed on a flow cytometer, either the BD FACSARIA for the screen or an Attune Nxt cytometer (ThermoFisher) for the follow-up experiments.

### Caspase activation assay

Cells were plated at 5,000 cells/well of a 384-well white opaque plates in corresponding cell growth media. Cells were treated with compounds at indicated concentrations for 6 hours (37 °C, 5% CO2) with a final DMSO concentration of 0.3%. Caspase-3/7 activation was subsequently determined using a Caspase-Glo 3/7 Reagent (Promega Corporation) as described in manufacturer’s instructions. Dose-response curves were plotted and analyzed (including EC50 determination) using GraphPad Prism. Percentage of caspase activation was calculated against the maximum caspase activation value (100%) obtained with a combination of MCL1, BCL2, and Bcl-xL inhibition.

### Cell viability assay

Cells were plated at 5,000 cells/well of a 384-well white opaque plates in corresponding cell growth media. Cells were treated with compounds at indicated concentrations for 24 hours (37 °C, 5% CO2) with a final DMSO concentration of 0.3% and assayed for viability using the CellTiter-Glo Reagent (Promega Corporation) as described in manufacturer’s instructions. Results were normalized to the samples without treatment at time 0. GI50values were calculated using nonlinear regression algorithms in GraphPad Prism.

### Immunoblotting

Primary antibodies against the following proteins were used: CUL5 (ab184177; Abcam); RNF7 (ab181986; Abcam); UBE2F (ab185234; Abcam); Noxa (OP180; Millipore); Bim (ab32158; Abcam); Bid (ab32060; Abcam); Puma (ab9643; Abcam); Bad (ab62465; Abcam); Bik (ab52182; Abcam); Beclin (ab207612; Abcam); Bmf (ab9655; Abcam); MCL1 (D35A5 #5453; CST); Bcl-xL (54H6; CST); c-PARP (19F4 #9546; CST) Ubiquitin (P4D1 #3936; CST); HA (C29F4; CST); GAPDH (14C10 #2118; CST); p53 (DO-1 sc-126; Santa Cruz Biotechnology); HSP90 (sc-69703; Santa Cruz Biotechnology). For each protein antibody, manufacturer’s recommended dilutions were used. Mouse or rabbit immunoglobulin G was visualized at a 1:10,000 dilution: donkey anti-mouse 680 (925-68022; LI-COR); donkey anti-rabbit 680 (925-68023; LI-COR); donkey anti-mouse 800 (925-32212; LI-COR); donkey anti-rabbit 800 (925-32213; LI-COR). Blots were imaged on an Odyssey CLx Imaging System (LI-COR).

### Immunoprecipitation

LK2 wild type and CUL5-KO cells were transfected with equal amounts of His-Ubiquitin and either HA-Noxa or HA-Bim. 48 hours after transfection, cells were treated with 100μM epoxomicin for 8 hours. Cells were harvested and lysed in either denaturing buffer (8M Urea, 300mM NaCl, 0.5% NP-40, 50mM Na2HPO4, 50mM Tris pH8.0) or lysis buffer from ThermoFisher (Pierce™ HA-Tag Magnetic IP/CoIP Kit #88838). Denatured extracts were bound to magnetic Ni-NTA beads (Qiagen #36111), while other lysed extracts were bound to magnetic anti-HA beads. Beads were washed 5 times and eluted in 1X Laemmli buffer by boiling for 5 minutes. Samples were loaded on a gel and processed for immunoblotting.

### siRNA treatment

Cells were reverse-transfected in 6-well plates using RNAiMAX (Thermo Fisher Scientific) with 50nM siRNA. 48h following siRNA treatment cells were treated for the Cell Event apoptosis assay as indicated and also harvested to verify knockdown by qRT-PCR. BCL2L11 (Bim) siRNA SMARTpool was from Dharmacon (L-004383-00-0005) and BAK siRNA was obtained from Ambion (Life Technologies 4457298).

---

**Figure S1.**
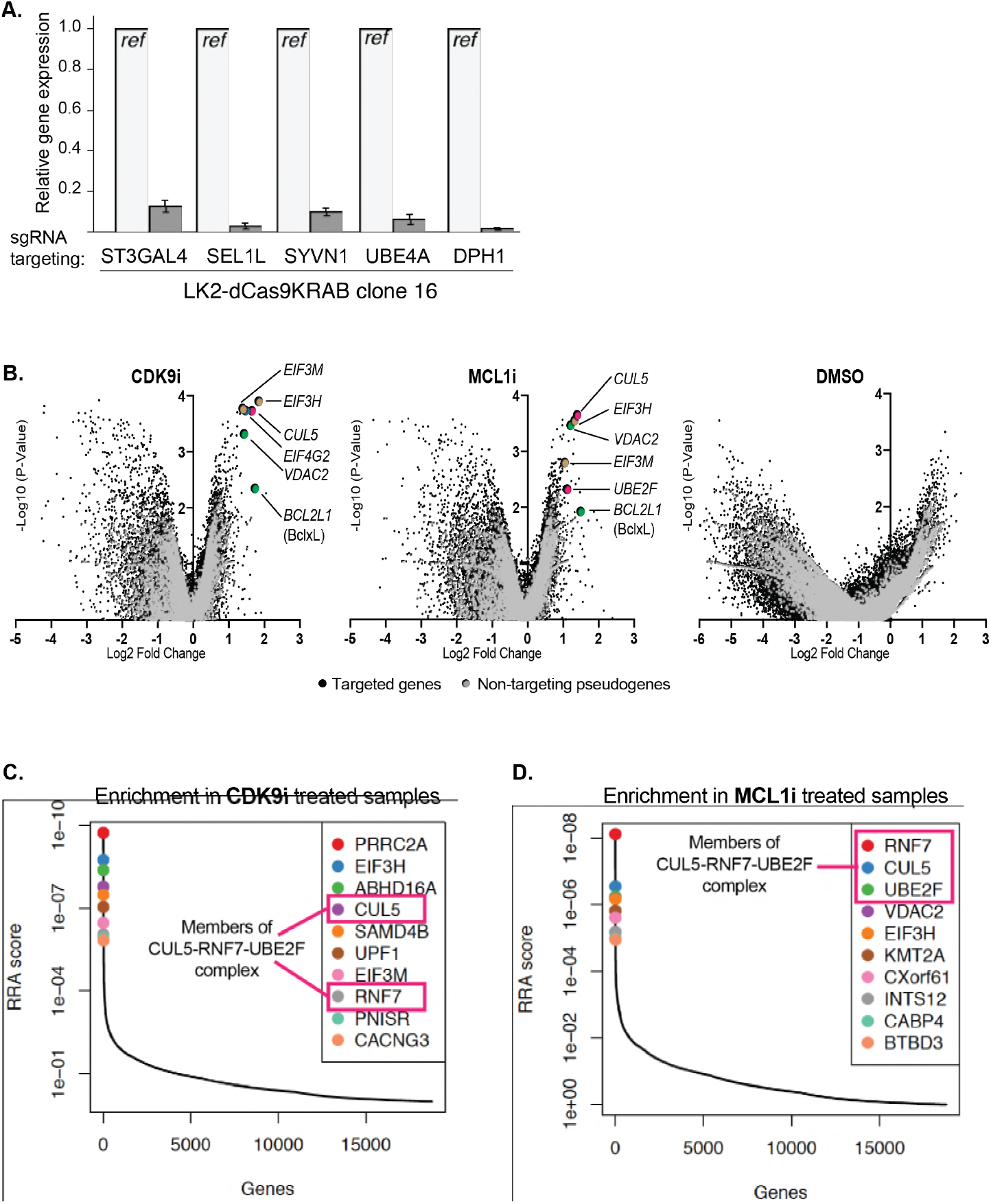
Performing the genome-wide CRISPRi screens and analysis of results. **(A)** Bar graph showing knockdown of five genes by qRT-PCR to confirm functionality of the dCas9-KRAB CRISPRi construct in LK2 cells. **(B)** Volcano plots (as in **Figure 2B**) showing significantly enriched and depleted genes when treated with CDK9i (left), MCL1 (middle) and DMSO (right). Black dots indicate targeted genes. Grey dots indicate negative control pseudogenes with non-targeting guides. Highlighted points of different colors show genes with significant fold changes with annotated roles in apoptosis (green), factors of the CUL5-RNF7-UBE2F ubiquitin complex (magenta), and members of the eIF3 complex (beige). **(C + D)** Relative sgRNA enrichment was also determined by the MAGeCK analysis pipeline. Top ten targeted genes following CDK9i and MCL1i treatment are shown in **(C)** and **(D)** respectively. sgRNAs targeting members of the CUL5-RNF7-UBE2F complex (outlined in magenta box) were once again identified as enriched and were amongst the most significant hits.

**Figure S2.**
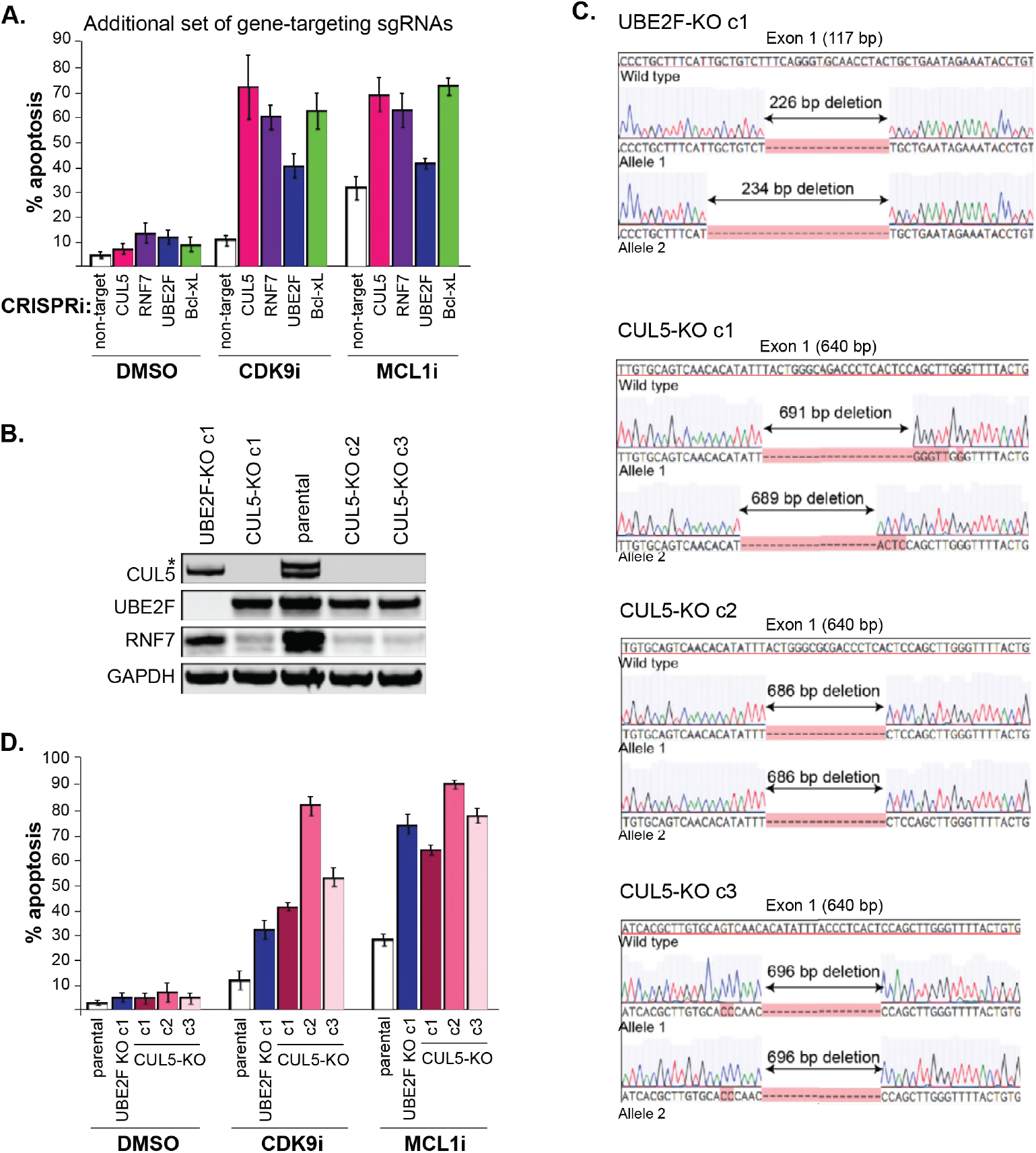
Generation of CUL5- and UBE2F-knockouts also resensitize cells to MCL1 inhibition. **(A)** Induction of apoptosis when cells are depleted of CUL5, RNF7, UBE2F or Bcl-xL with an additional set of sgRNAs than those used in **Figure 3D** and are treated with CDK9i (3μM for 6h), MCL1i (1μM for 12h) or DMSO. Percentage of apoptosis determined by flow cytometry for detection of the fluorogenic apoptotic detection reagent Cell Event. Error bars are standard deviations of three biological replicates. **(B)** Western blotting validating loss of CUL5 and UBE2F protein in the three CUL5-KO clones and one UBE2F-KO clone respectively. As shown before, protein levels of RNF7 are also reduced when CUL5 is absent. GAPDH, loading control. **(C)** Sanger sequencing traces of the alleles generated in the CUL5- and UBE2F knockouts. Exon 1 size is indicated, and resulting deletions are shown. **(D)** CUL5-KO and UBE2F-KO clones were treated with 3μM CDK9i for 6 hours or 1μM MCL1i for 12 hours. Percent apoptosis determined using Cell Event detection by flow cytometry. Error bars derived from standard deviations of three biological replicates.

**Figure S3.**
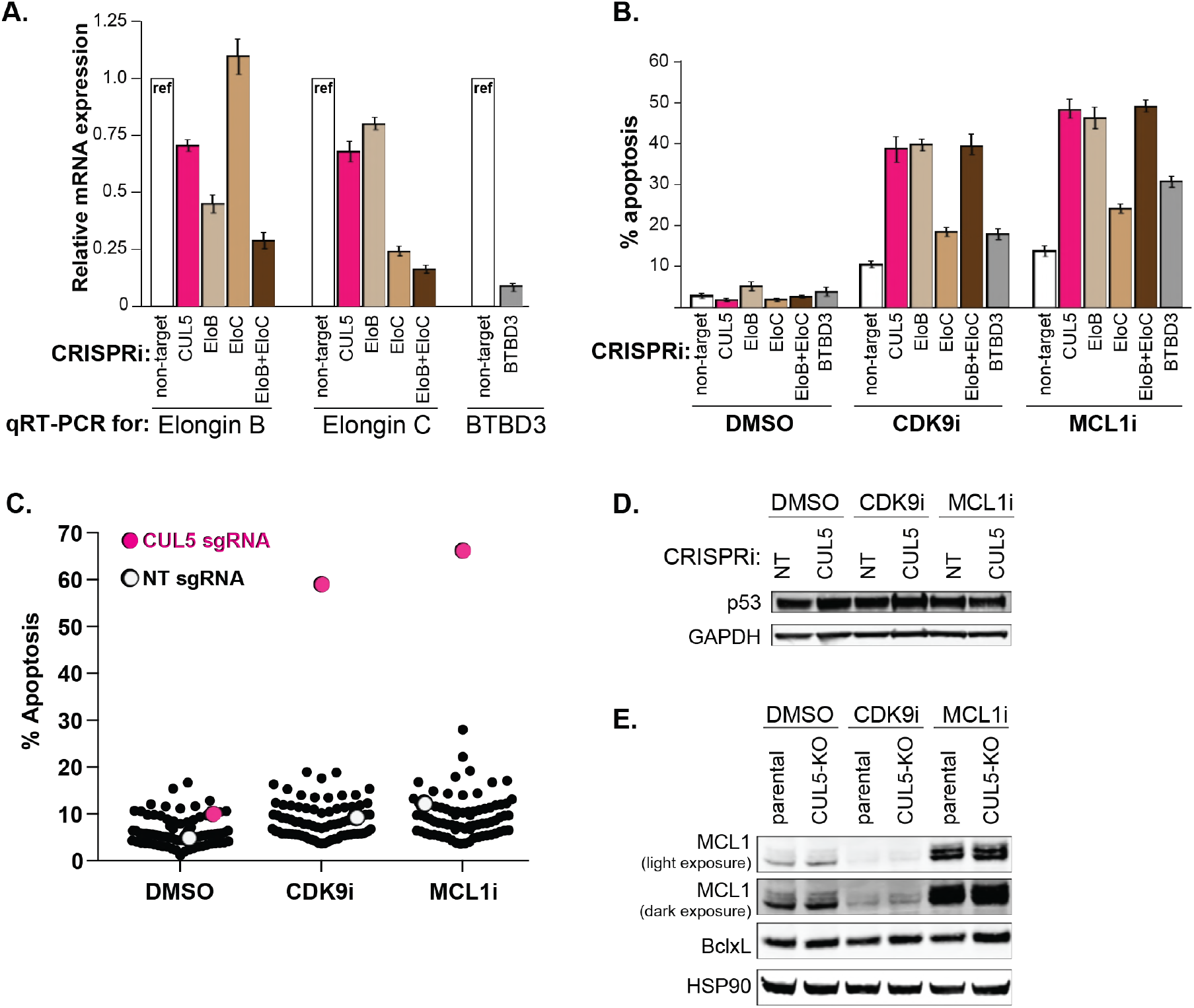
Knockdown of Elongin B but not Elongin C or BTBD3 also resensitizes cells to MCL1 inhibition. **(A)** qRT-PCR to confirm efficient knockdown of Elongin B, Elongin C and BTBD3. **(B)** Knockdown of Elongin B (EloB), but not Elongin C (EloC), also elicits similar levels of apoptosis as compared to CUL5-knockdown when treated with CDK9i or MCL1i. Knockdown of BTBD3 in conjunction with CDK9i and MCL1i has a modest increase in apoptosis. Apoptosis measured by Cell Event, error bars are standard deviations of three biological replicates. **(C)** Plotted are percentages of apoptosis as determined by flow cytometry measuring Cell Event fluorescence after 3μM drug treatment for 6 hours, as indicated on the X axis. Each dot corresponds to an individual CRISPRi gene knockdown. Non-target (NT) sgRNA serves as the negative control (white) and CUL5 knockdown serves as the positive control (magenta). **(D)** Western blot for p53 in NT and CUL5-depleted cells treated with DMSO, CDK9i (3μM for 6h) or MCL1i (1μM for 12h). GAPDH serves as loading control. **(E)** Western blot of MCL1 and Bcl-xL in LK2 and CUL5-KO c1 cells treated with DMSO, CDK9i or MCL1i. HSP90 serves as loading control.

**Figure S4.**
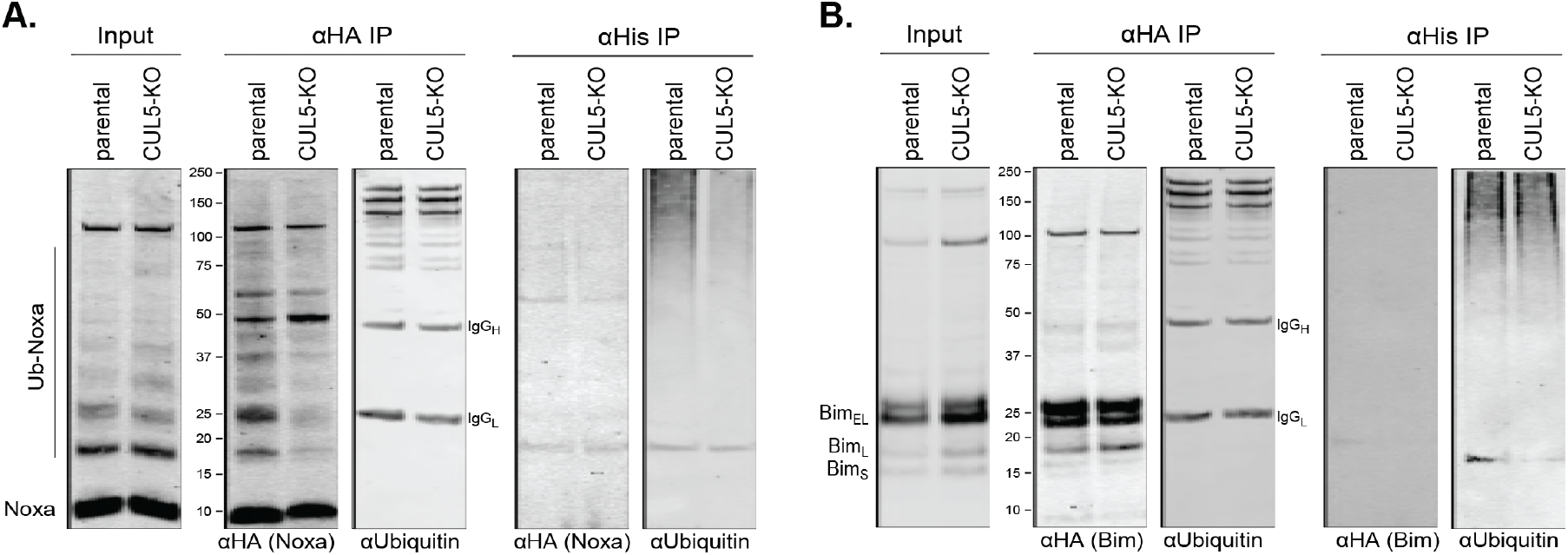
Indirect regulation of Bim and Noxa by the CUL5-RNF7-UBE2F complex. **(A)** LK2 and CUL5-KO c1 cells were transiently transfected with HA-Noxa and His-Ubiquitin. 48 hours following transfection cells were incubated with 100μM epoxomicin for 8 hours and subsequently harvested for immunoprecipitations (IP) with anti-HA or Ni-NTA. Western blot show 1% input and 50% of IP loaded and probed with the indicated antibodies. **(B)** LK2 and CUL5-KO c1 cells were transiently transfected with HA-Bim and His-Ubiquitin. 48 hours following transfection cells were incubated with 100μM epoxomicin for 8 hours and subsequently harvested for immunoprecipitations (IP) with anti-HA or Ni-NTA. Western blot show 1% input and 50% of IP loaded and probed with the indicated antibodies.

**Figure S5.**
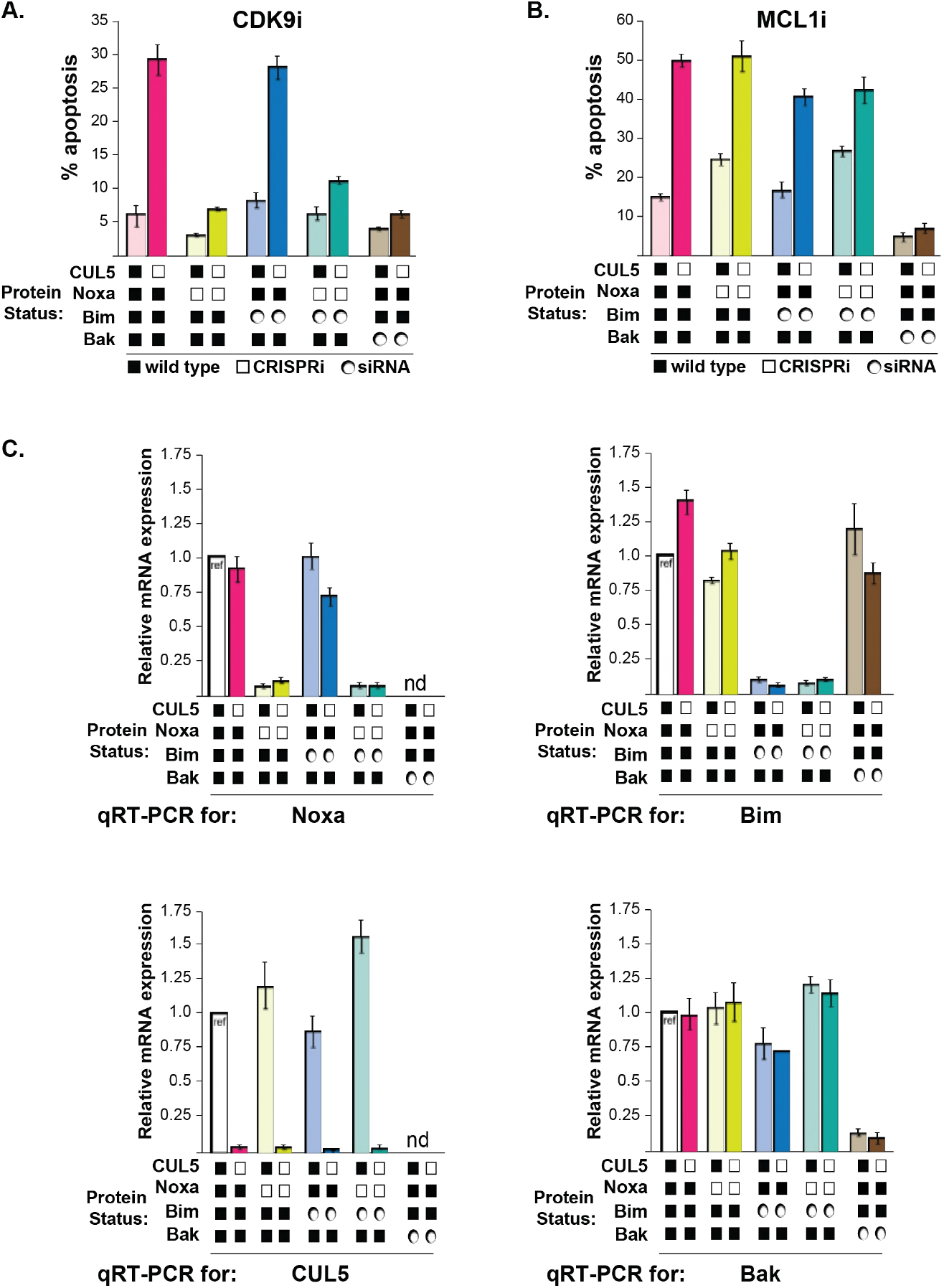
CUL5-kd cells regain resistance to CDK9i when Noxa is depleted. **(A)** As in Figure 5, CUL5 and Noxa were knocked down in dCas9-KRAB-expressing LK2 cells by selecting for corresponding sgRNAs. Bim and Bak were knocked down by siRNA. Apoptosis was measured using Cell Event detection by flow cytometry after treatment with 3μM CDK9i for 6 hours. Bar graphs depict an independent set of transductions and siRNA transfections from **Figure 5**. Error bars are standard deviations of 3 biological drug treatment replicates. **(B)** Genetic manipulations were performed and apoptosis was measured as in **(A)** following MCL1i treatment. **(C)** Representative data set illustrating knockdown efficiency by qRT-PCR for genetic manipulations shown in (**Figure 5A-B**). Error bars are standard deviations of technical replicates. qRT-PCR was conducted to validate knockdown for each biological replicate of sgRNA transduction and siRNA transfection. nd, not determined.

**Table S1 (related to Figure 1).** A subset of resistant and sensitive NSCLC lines. For each cell line, the table indicates *MCL1* copy number, MCL1:Bcl-xL protein ratio and EC_50_ concentrations for both CDK9i and MCL1 treatments.

|  | MCL1 Copy Number | Mcl1:BclxL Protein Ratio | MCL1i GI<sub>50</sub> (μM) | MCL1i Caspase EC<sub>50</sub> (μM) | CDK9i Caspase EC<sub>50</sub> (μM) |
| --- | --- | --- | --- | --- | --- |
| SKLU1 | 1.5 | 0.41 | 10.000 | 30.000 | 30.000 |
| HCC827 | 2.3 | 4.38 | 10.000 | 30.000 | 30.000 |
| H460 | 3.2 | 16.58 | 10.000 | 30.000 | 30.000 |
| H1734 | 3.6 | 2.63 | 6.150 | 20.056 | 30.000 |
| LK2 | 4.7 | 3.71 | 7.040 | 30.000 | 30.000 |
| H1395 | 6.6 | 3.79 | 10.000 | 15.404 | 30.000 |
| H23 | 3.1 | 39.59 | 0.383 | 0.198 | 0.183 |
| H2110 | 3.9 | 18.29 | 0.113 | 0.226 | 0.270 |
| H1568 | 6.5 | 14.26 | 0.111 | 0.3 | 0.151 |

## SUPPLIED AS ADDITIONAL SUPPORTING DATA

**Table S2 (related to Figure 2). ScreenProcessing: sgRNA counts**.

**Table S3 (related to Figure 2). ScreenProcessing: sgRNA phenotype scores**.

**Table S4 (related to Figure 2). ScreenProcessing: gene phenotype scores**.

**Table S5 (related to Figure S1). MAGeCK: sgRNA summary for CDK9i**.

**Table S6 (related to Figure S1). MAGeCK: gene summary for CDK9i**.

CDK9i treatment condition compared to ‘background’. Ordered by top enriched genes.

**Table S7 (related to Figure S1). MAGeCK: sgRNA summary for MCL1i**.

**Table S8 (related to Figure S1). MAGeCK: sgRNA summary for MCL1i**.

MCL1i treatment condition compared to background. Ordered by top enriched genes.

**Table S9 (related to Figure S4). Analysis of apoptosis following knockdown of putative CUL5 substrate adaptors challenged with CDK9i or MCL1i**.

**Table S10 (related to Figure 3 and S3). Protospacer sequences**.

**Table S11. Primer Sequences**

